# Within-Host Diversity and Vertical Transmission of Group B *Streptococcus* Among Mother-infant Dyads in The Gambia

**DOI:** 10.1101/760512

**Authors:** Ebenezer Foster-Nyarko, Madikay Senghore, Brenda A. Kwambana-Adams, Nabil-Fareed Alikhan, Anuradha Ravi, James Jafali, Kaddijatou Jawneh, Amara Jah, Maimuna Jarju, Fatima Ceesay, Sainabou Bojang, Archibald Worwui, Aderonke Odutola, Ezra Ogundare, Mark J. Pallen, Martin Ota, Martin Antonio

## Abstract

**Introduction:** Understanding mother-to-infant transmission of Group B *Streptococcus* (GBS) is vital to the prevention and control of GBS disease. We investigated the transmission and phylogenetic relationships of mothers colonised by GBS and their infants in a peri-urban setting in The Gambia.

**Methods:** We collected nasopharyngeal swabs from 35 mother-infant dyads at weekly intervals from birth until six weeks post-partum. GBS was isolated by conventional microbiology techniques. Whole-genome sequencing was performed on GBS isolates from one mother-infant dyad (dyad 17).

**Results:** We recovered 85 GBS isolates from the 245 nasopharyngeal swabs. GBS was isolated from 16.33% and 18.37% of sampled mothers and infants, respectively. In 87% of cultured swabs, the culture status of an infant agreed with that of the mother (Kappa p-value <0.001). In dyad 17, phylogenetic analysis revealed within-host strain diversity in the mother and clone to her infant.

**Conclusion:** GBS colonisation in mothers presents a significant risk of colonisation in their infants. We confirm vertical transmission from mother to child in dyad 17, accompanied by within-host diversity.

## Introduction

*Streptococcus agalactiae*, also known as Group B *Streptococcus* (GBS), is notable for causing meningitis, sepsis and death in young infants across the globe [1]. A recent global estimate reported that, in 2015, GBS caused 90,000 deaths in infants aged younger than three months of age [1]; Africa accounted for 54% of estimated cases and 65% of all foetal/infant deaths. Maternal colonisation of GBS is an important source of infection in new-borns—up to 70% of infants born to colonised mothers are likely to be colonised themselves [2]. However, the microbial population genetics of mother-infant transmission of GBS remains poorly understood.

Previous attempts to investigate the transmission of GBS have relied on mother-child colonisation status and typing methods with limited resolution, such as serotyping and multi-locus sequence typing (MLST) [3–5]. However, whole-genome sequencing studies on other pathogens have shown that a cloud of genomic diversity—encompassing multiple single-nucleotide variants and arising from micro-evolutionary events—can exist within populations that appear homogenous by conventional typing approaches [6–9]. In particular, application of next-generation sequencing technology to multiple colonies from the same individual, whether from single or multiple samples, can shed light on the influence of within-host diversity on chains of transmission [6–11]. Intervention and prevention strategies targeted at GBS disease, such as maternal immunisation, would be enhanced by an improved understanding of the within-host diversity associated with this important pathogen.

Here, we combine conventional microbiologic investigations with whole-genome sequencing to elucidate mother-infant transmission of GBS among a cohort of mother-infant dyads in a peri-urban setting in The Gambia.

## Methods

### Study design and study participants

We carried out a prospective, longitudinal study of mother-infant dyads. Participants were recruited at the Fajikunda Health Centre, which is located in a peri-urban setting in the coastal region of The Gambia, about 16km from the capital, Banjul. The Fajikunda health centre and the Fajikunda community have been described elsewhere [12]. Mothers aged 18 years and older attending the study clinic were consented before delivery and assigned study numbers. The inclusion criteria included 1) willingness to deliver at the health centre; 2) absence of pregnancy-related complications; and 3) availability post-delivery for the duration of sampling (two months). Trained fieldworkers obtained written informed consent and administered questionnaires to study participants. History of antibiotic use two weeks prior to enrolment, age and area of settlement were recorded. Recruitment was done between August 2011 and November 2012. The mother-child dyads were swabbed consecutively at seven weekly time points (designated Week1-Week7) from birth till two months of age.

### Ethical approval

The study was approved by the Joint Gambia Government and the Medical Research Council Unit The Gambia at London School of Hygiene and Tropical Medicine (MRCG at LSHTM) Ethics Committee (study number SCC 1173).

### Sample collection

Nasopharyngeal swabs were collected from participants at birth and once weekly for the subsequent six weeks. Trained nurses collected the samples as described previously [12]. The swabs were immediately immersed into Skim-milk Tryptone Glucose Glycerol broth, kept in cooler boxes at 2-8°C and transported to the MRCG at LSHTM’s laboratories at Fajara within six hours. Upon receipt in the laboratory, the samples were homogenised by vortex for a minimum of 20 seconds and stored at −80°C until further processing.

### Culture and isolation of GBS

Aliquots of the nasopharyngeal swab suspensions were enriched in Todd-Hewitt Broth with 5 % Yeast Extract (Oxoid, Basingstoke, UK) and rabbit serum (B&K Universal Ltd, Grimston, East Yorkshire, UK) prior to culture, as described previously [13]. Next, the nasopharyngeal swab suspensions were plated on 5% Sheep blood agar supplemented with crystal violet.

GBS isolates were identified using conventional methods. Briefly, candidate bacterial colonies demonstrating beta-haemolytic activity on blood agar, bacitracin negative, catalase negative, Christie, Atkins, Munch and Peterson (CAMP) test positive were confirmed as GBS using the Streptex® Streptococcal grouping kit (Remel & Oxoid, Thermo Fisher Scientific, Basingstoke, UK). Single colonies were purified and stored from each nasopharyngeal swab culture. All strains were stored in 20% glycerol broth at −80°C until further processing.

### Genomic DNA extraction and sequencing

GBS isolates from one informative mother-child dyad (dyad 17) was chosen for whole-genome sequencing. To provide a context for cluster analysis of the isolates, we included an isolate each from Mother 04 (Mother4_Week4), Child 19 (Child19_Week4) and Child 07 (Child07_Week4). Selected GBS isolates were streaked on 5% sheep blood agar plates and incubated overnight in ambient air. Following overnight incubation, a single colony from each plate was picked, inoculated into 1ml Luria-Bertani broth and incubated for 4 hours at 37°C. The Luria-Bertani broth cultures were then spun at 3500rpm for two minutes and lysed using a cocktail comprised of lysozyme, proteinase K, 10%SDS and RNase A in Tris EDTA buffer (pH 8.0). The suspension was then placed on a thermomixer with vigorous shaking at 1600rpm; first at 37°C for 25 minutes and subsequently at 65°C for 15 minutes.

The DNA was extracted using solid-phase reversible immobilisation magnetic beads and precipitated with ethanol. Elution of the DNA was done in Tris-Cl. The quality of the DNA was assessed on the Thermo Scientific NanoDrop 2000 Spectrophotometer (Fisher Scientific UK Ltd, Loughborough, UK) using the A260/A280 and A260/A230 ratios to detect contaminating proteins and RNA respectively. The DNA concentrations were quantified using the Qubit HS DNA assay (Invitrogen) and normalised to 0.5ng/μL before preparation of sequencing libraries. The eluted DNA was stored at −20°C. Whole-genome sequencing was performed on the Illumina NextSeq 500 platform, following the Nextera XT DNA library preparation kit protocol (Illumina, San Diego, CA) using a mid-output kit.

### Bioinformatics analysis

The sequencing data was analysed on the Cloud Infrastructure for Microbial Bioinformatics platform [14]. The paired-end sequences were merged to enable error correction and accurate read coverage determination. The merged reads were then quality-checked using FastQC v0.11.7 (http://www.bioinformatics.babraham.ac.uk/projects/fastqc/). To determine the closest NCBI RefSeq genomes to our stains, we used RefSeqMasher (https://github.com/phac-nml/refseq_masher/tree/master/refseq_masher). The SGBS021 genome (RefSeq accession NZ_AUWC 00000000.1) was chosen as the closest match to our genomes. The reads were then assembled using Shovill (https://github.com/tseemann/shovill). We assessed the quality of assemblies using QUAST v 5.0.0, de6973bb (https://github.com/ablab/quast). Multi-locus sequence types were called from the assembled contigs. Annotation was done using Prokka [15]. We used Snippy v4.3.2 (https://github.com/tseemann/snippy) for variant calling and core gene alignment, and snp-dists v0.6 (https://github.com/tseemann/snp-dists) to compute Single Nucleotide Polymorphism (SNP) distances between the genomes. RAxML v8.2.4 [16] was used for maximum-likelihood phylogenetic inference based on a general time-reversible nucleotide substitution model with 1,000 bootstrap replicates. We used the GrapeTree MSTree V2 algorithm [17] to visualise and annotate phylogenetic trees. We used a whole-genome-sequence-based method for deducing GBS serotypes, “GBS_serotyper.pl” to detect the serotypes from the sequence reads as described elsewhere [18]. Briefly, this tool relies on SRST2 and the GBS pubmlst database (http://pubmlst.org/sagalactiae/) to predict serotypes from query DNA sequences.

### Data Management and Statistical analysis

Data were managed on an SQL integrated database with a Microsoft Access front-end linked to the demographic, clinical and laboratory databases. We used Stata version 12 (Stata Corp, College Station, TX) and R software (https://www.r-project.org) packages for the statistical analyses. Descriptive statistics were computed including frequency counts (n) and percentages (%), factor variables and mean (SD) or median (IQR) (normally or non-normally distributed continuous variables, respectively). To assess the agreement of GBS carriage between the mother and child pairs, McNemmar’s test and Weighted kappa coefficients for correlation of two raters (irr package) were applied. P values of less than 5% were considered significant to reject the null hypothesis.

## Results

### GBS colonisation

We recruited 35 mother-infant dyads (17 male infants; 18 females) from villages around the Fajikunda Health Centre in The Gambia (Table 1) and collected 245 nasopharyngeal swabs over a 7-week sampling period (seven swabs per study participant). None of the study participants reported antibiotic use in the two weeks prior to delivery. Microbiological culture recovered 85 GBS isolates.

**Table 1:** Strains used in this study

| **Study ID** | **Lab ID** |
| --- | --- |
| 1 | 001 Mother/Child |
| 2 | 004 Mother/Child |
| 3 | 005 Mother/Child |
| 4 | 006 Mother/Child |
| 5 | 007 Mother/Child |
| 6 | 009 Mother/Child |
| 7 | 010 Mother/Child |
| 8 | 011 Mother/Child |
| 9 | 012 Mother/Child |
| 10 | 013 Mother/Child |
| 11 | 014 Mother/Child |
| 12 | 015 Mother/Child |
| 13 | 016 Mother/Child |
| 14 | 017 Mother/Child |
| 15 | 018 Mother/Child |
| 16 | 019 Mother/Child |
| 17 | 020 Mother/Child |
| 18 | 021 Mother/Child |
| 19 | 022 Mother/Child |
| 20 | 023 Mother/Child |
| 21 | 027 Mother/Child |
| 22 | 028 Mother/Child |
| 23 | 029 Mother/Child |
| 24 | 031 Mother/Child |
| 25 | 032 Mother/Child |
| 26 | 035 Mother/Child |
| 27 | 036 Mother/Child |
| 28 | 037 Mother/Child |
| 29 | 040 Mother/Child |
| 30 | 044 Mother/Child |
| 31 | 045 Mother/Child |
| 32 | 046 Mother/Child |
| 33 | 047 Mother/Child |
| 34 | 048 Mother/Child |
| 35 | 049 Mother/Child |

GBS was found in 16.33% (40/245) in the mothers and in 18.37% (45/245) of infants (Table 2). In 87.35% (214/245) of cultured swabs, the culture status of an infant (i.e. positive or negative for GBS) agreed with that of the mother at a given time point (Kappa *p-*value <0.001) (Table 3). GBS positivity among the mother-infant dyads across the six weeks of sampling is shown in Figure 1.

**Table 2:** Prevalence of Group B *streptococcus* across time

| Visit<sup>∞</sup> | N | Mother |  | Child |  |
| --- | --- | --- | --- | --- | --- |
|  |  | n | % | n | % |
| 1 | 35 | 6 | 17.14% | 6 | 17.14% |
| 2 | 35 | 7 | 20% | 9 | 25.71% |
| 3 | 35 | 9 | 25.71% | 10 | 28.57% |
| 4 | 35 | 8 | 22.86% | 7 | 20% |
| 5 | 35 | 5 | 14.29% | 6 | 17.14% |
| 6 | 35 | 2 | 5.71% | 3 | 8.57% |
| 7 | 35 | 3 | 8.57% | 4 | 11.43% |
| **Total** | **245** | **40** | **16.33%** | **45** | **18.37%** |

<sup>∞</sup>Visit 1 represents sampling at birth; visits 2-7 are weekly sampling points from week 1 to week 6.

**Table 3:** Measure of Agreement between mother and child

| **Visit<sup>∞</sup>** | **Observed** |  | **Kappa's P value** |
| --- | --- | --- | --- |
|  | **n/N** | **Percent** |  |
| 1 | 31/35 | 88.57% | <0.001 |
| 2 | 27/35 | 77.14% | 0.0334 |
| 3 | 28/35 | 80.00% | 0.0033 |
| 4 | 30/35 | 85.71% | <0.001 |
| 5 | 34/35 | 97.14% | <0.001 |
| 6 | 32/35 | 91.43% | 0.0311 |
| 7 | 32/35 | 91.43% | 0.0017 |
| Total | 214/245 | 87.35% | <0.001 |

<sup>∞</sup>Visit 1 represents sampling at birth; visits 2-7 are weekly sampling points from week 1 to week 6.

**Figure 1:**
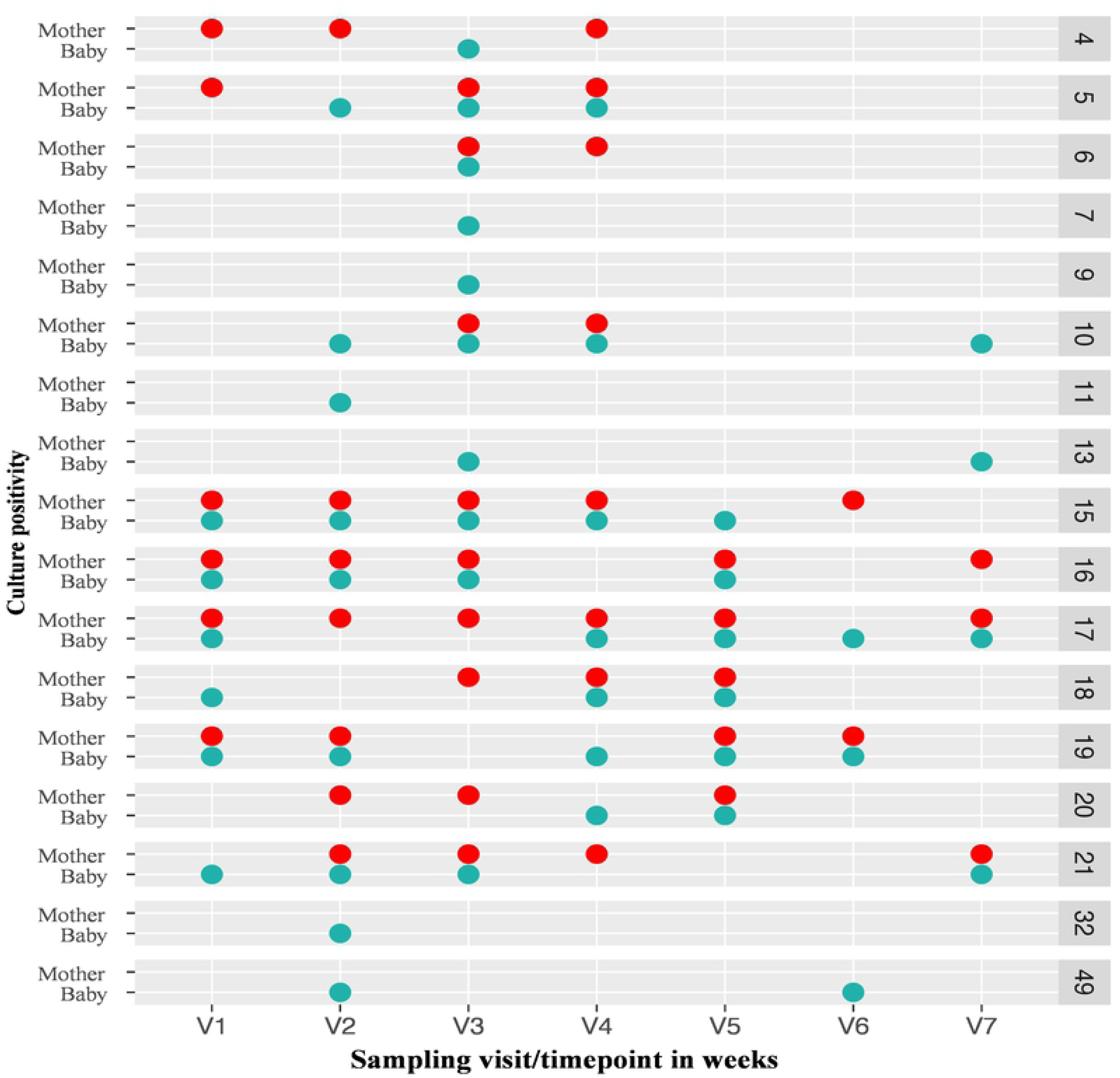
Temporal colonisation of GBS among the Study participants. V1, birth; V2-V7, weeks 1 to 6. The lab IDs of mother-infant pairs are shown on the secondary y-axis (4, 5, 17 etc). Red, mother; light sea green, baby. Of six infants who were colonised by GBS at birth, four were born to colonised mothers (dyads 15, 16, 17 and 19). Of the GBS colonisations detected post-birth, seven were found in infants alone or prior to the isolation of GBS from their mothers (dyads 7, 9, 10, 11, 13, 32 and 49). In the remaining instances, GBS was first isolated from both mother and infant at the third timepoint (dyad 6, week 2) or from the mother prior to isolation from the infant (dyad 20).

Four mother-infant dyads had GBS isolated from both mother and child at birth (dyads 15, 16, 17 and 19). Of the remaining four instances where an isolate was recovered at birth, two were in mothers (mothers 4 and 5) only and the other two in infants only (infants 18 and 21). In seven out of nine instances where GBS was isolated post-birth, the infant was found to have GBS when the mother was negative or before the isolation of GBS from the mother (dyads 7, 9, 10, 11, 13, 32 and 49). In the remaining instances, GBS was first isolated from both mother and infant at week 2 (dyad 6) or from the mother two weeks prior to isolation from the infant (dyad 20).

Only dyad 17 had an isolate recovered at each weekly sampling visit. Eleven genomes in all were recovered from dyad 17; however, one isolate was lost to contamination, leaving ten genomes for genomic analysis. All ten isolates from dyad 17 belonged to ST196 and were assigned by in-silico capsular typing to serotype II. GBS isolates Child07_Week3 and Mother04_Week4 were both assigned to ST10 and serotype II; Child19_Week4 was assigned to serotype V and ST26.

### Phylogenomic diversity among isolates from dyad 17

A phylogenetic tree constructed from core-genome SNPs shows that strains from mother and infant 17 cluster together and are separated from the SGBS021 genomic reference and our other three GBS genomes (Figure 2). Genomes from dyad 17 were at most 7 SNPs apart, whereas the next closely related genome was 1,2447 SNPs apart (Table 4). GBS genomes recovered from mother 17 at weeks one and week four and all the genomes recovered from infant 17 were identical (Table 4). However, we observed a cloud of minor SNP variants within the isolates from mother 17, encompassing four genotypes. Genotype-1 (from Week1 and Week4) differed from the genotype-2 recovered at week two by one SNP. Genotype-2 differed from genotype-3 recovered at week three by seven SNPs; genotype-3 was one SNP apart from genotype-4 recovered at Week5.

**Table 4:** Single Nucleotide Polymorphism (SNP) matrix showing the SNP diversity among the study isolates

| Strain | Mother04_Week3 | Child07_Week2 | Mother17_Week1 | Mother17_Week2 | Mother17_Week3 | Mother17_Week4 | Mother17_Week6 | Child17_Birth | Child17_Week3 | Child17_Week4 | Child17_Week5 | Child17_Week6 | Child19_Week3 | Reference |
| --- | --- | --- | --- | --- | --- | --- | --- | --- | --- | --- | --- | --- | --- | --- |
| Mother04_Week3 | 0 | 46 | 1677 | 1678 | 1683 | 1677 | 1683 | 1677 | 1677 | 1677 | 1677 | 1677 | 12554 | 3308 |
| Child07_Week2 | 46 | 0 | 1723 | 1724 | 1729 | 1723 | 1729 | 1723 | 1723 | 1723 | 1723 | 1723 | 12594 | 3354 |
| Mother17_Week1 | 1677 | 1723 | 0 | 1 | 6 | 0 | 6 | 0 | 0 | 0 | 0 | 0 | 12445 | 3023 |
| Mother17_Week2 | 1678 | 1724 | 1 | 0 | 7 | 1 | 7 | 1 | 1 | 1 | 1 | 1 | 12446 | 3024 |
| Mother17_Week3 | 1683 | 1729 | 6 | 7 | 0 | 6 | 1 | 6 | 6 | 6 | 6 | 6 | 12439 | 3017 |
| Mother17_Week4 | 1677 | 1723 | 0 | 1 | 6 | 0 | 6 | 0 | 0 | 0 | 0 | 0 | 12445 | 3023 |
| Mother17_Week6 | 1683 | 1729 | 6 | 7 | 1 | 6 | 0 | 6 | 6 | 6 | 6 | 6 | 12440 | 3018 |
| Child17_Birth | 1677 | 1723 | 0 | 1 | 6 | 0 | 6 | 0 | 0 | 0 | 0 | 0 | 12445 | 3023 |
| Child17_Week3 | 1677 | 1723 | 0 | 1 | 6 | 0 | 6 | 0 | 0 | 0 | 0 | 0 | 12445 | 3023 |
| Child17_Week4 | 1677 | 1723 | 0 | 1 | 6 | 0 | 6 | 0 | 0 | 0 | 0 | 0 | 12445 | 3023 |
| Child17_Week5 | 1677 | 1723 | 0 | 1 | 6 | 0 | 6 | 0 | 0 | 0 | 0 | 0 | 12445 | 3023 |
| Child17_Week6 | 1677 | 1723 | 0 | 1 | 6 | 0 | 6 | 0 | 0 | 0 | 0 | 0 | 12445 | 3023 |
| Child19_Week3 | 12554 | 12594 | 12445 | 12446 | 12439 | 12445 | 12440 | 12445 | 12445 | 12445 | 12445 | 12445 | 0 | 12277 |
| SGBS021 Reference genome | 3308 | 3354 | 3023 | 3024 | 3017 | 3023 | 3018 | 3023 | 3023 | 3023 | 3023 | 3023 | 12277 | 0 |

3 Pair-wise single nucleotide polymorphism (SNP) comparison of study genomes, showing that genomes from dyad 17 were 0-7 SNPs apart, and
4 1,2447 SNPs apart from the next closest relative (Table 4). Similarly, genomes from mother 03, child 07 and child 19 were 46-12594 SNPS 5 apart from each other. The GBS genomes recovered baby 17 were identical to each other, and to that recovered from mother 17 at weeks one and 6 week four. Four minor SNP variants were observed among the genomes from mother 17. Genotype-1 (Week1 and Week4) differed from the 7 genotype-2 (Week2) by one SNP. Genotype-2 differed from genotype-3 (Week3) by seven SNPs; genotype-3 was one SNP apart from genotype- 8 4 (Week5).

**Figure 2:**
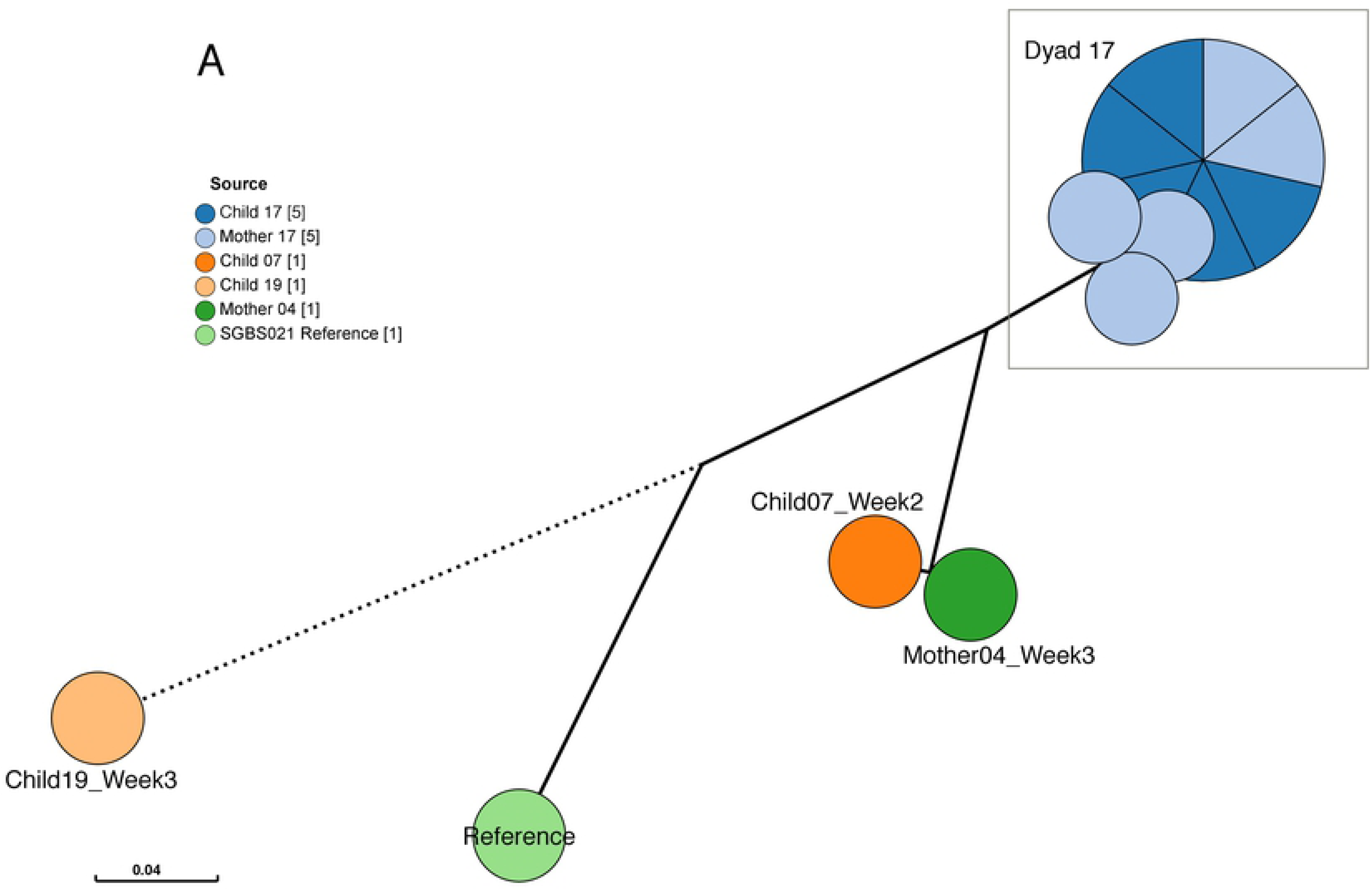
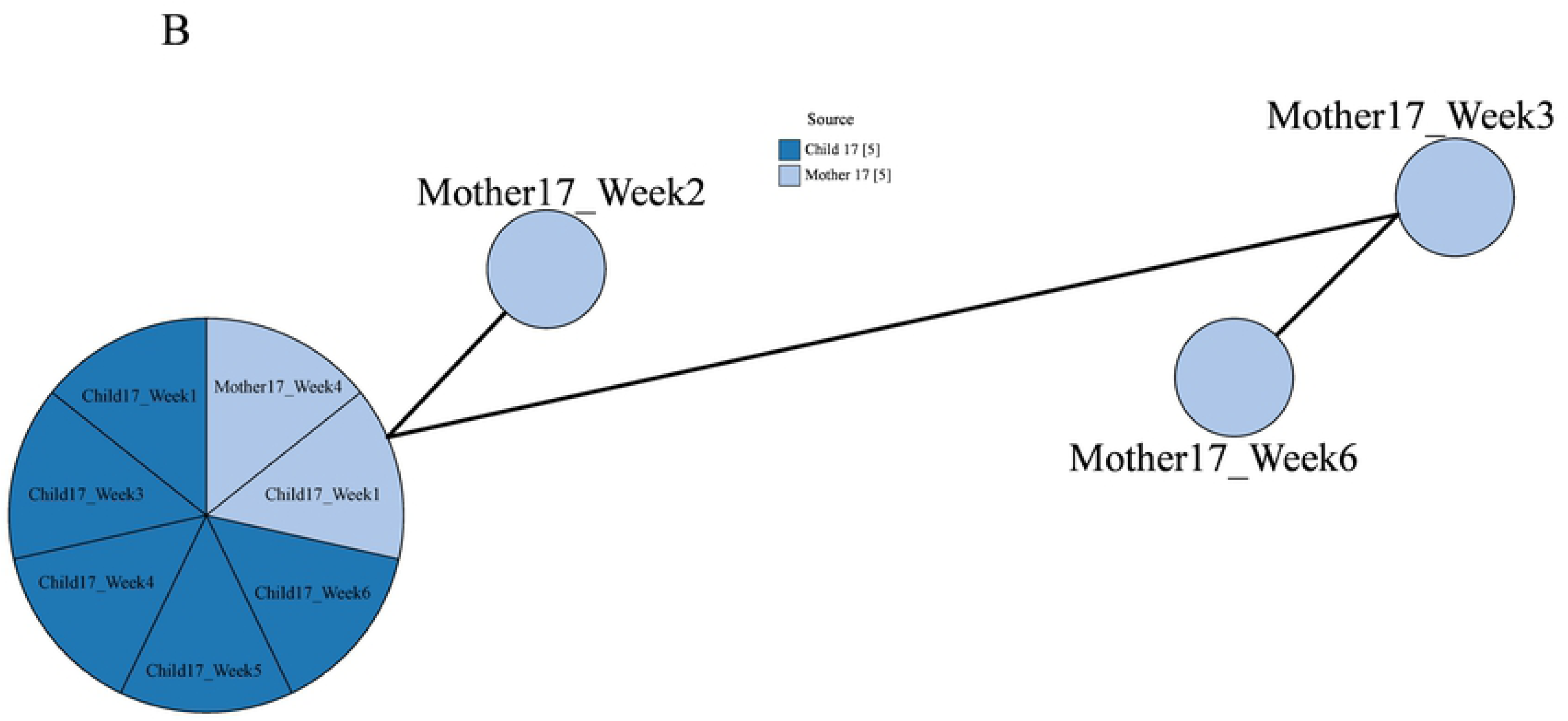
A&B: Phylogenetic relationship between GBS isolates from mother-infant dyad 17 and three other genomes (2 mothers and two infants) The genomes are labelled with the dyad number from which they were derived. Mother17 & Child17 represent genomes from dyad17. Figure 2A shows the genomes from dyad 17 cluster together and are several 1000s of SNPS apart from the genomic reference and the other genomes, indicating that the genomes from mother 17 and infant 17 from the same genetic pool (See table 6). The genomes derived from infant 17 were identical to each other and to the maternally-derived genomes at weeks one (Mother17_Week1) and four (Mother17_Week4). Also, the non-dyad 17 genomes have at least 46 SNPs between them (see Table 4). Branches longer than 0.2 were shortened (denoted by broken lines). Figure 2B shows the phylogenetic relationship between genomes within dyad 17. Not shown here are the number of SNPS between the respective genomes within dyad 17. Among the genomes from infant 17, Child17_Week5 and Child17_Week6 are the closest, separated by 3SNPS. Of the maternal genomes, Mother17_Week1 and Mother17_Week2 are the closest, with 5 SNPs separating them. The highest pairwise distance between the genomes within dyad 17 is 25SNPs (between Mother17_Week3 and Mother17_Week6).

## Discussion

Our results highlight the utility of nasopharyngeal sampling in the investigation of GBS colonisation and transmission. We found a nasopharyngeal colonisation prevalence of 16.33% in mothers and 18.37% in their infants. These results are consistent with a recent meta-analysis of global maternal GBS colonisation, which reported a prevalence of 15% regardless of sampling site [19], and with the neonatal prevalence rate we previously reported in a two-month-old infant cohort [13].

We have documented vertical transmission followed by persistent colonisation of an ST196 GBS strain in a mother-infant dyad. ST196 is a successful coloniser [3] as well as an essential cause of invasive disease in adults [20]. This clone has also been linked to transmission between humans and cattle [21]. Several studies have associated ST196 with serotype IV [22]. We are aware of only one other study which reported one strain out of 102 prospectively collected strains to be ST196 with a serotype II capsular backbone [23]. Diversity among the maternal ST196 genomes tended to increase across time, with microevolution producing a cloud of closely related genetic variants. Four distinct GBS genotypes were recovered from mother 17; however, all the GBS genomes from infant 17 were identical to each other and to the maternal GBS genomes recovered from weeks one and four, suggesting that a single genotype was transmitted to the infant at birth and persisted during the sampling window.

Of nine instances where GBs was detected post-birth, seven were found in the infants alone or prior to detection in their mothers. This finding may be due to the insensitivity of the culture method, or it may affirm the role of non-maternal sources in neonatal GBS infection. The importance of horizontal transmission—from healthcare providers, family and friends— in the investigation of GBS infection in new-borns has been documented [24–26]. Enhanced surveillance of the burden of horizontal transmission is warranted to inform early intervention and prevention strategies.

## Conclusion

Although we sequenced a limited number of GBS genomes, our findings provide valuable insight into the dynamics of transmission in our setting. Future longitudinal studies incorporating large-scale whole-genome sequencing of multiple isolates from mother-infant dyads will further enhance our understanding of the full scope of the genomic diversity of colonising GBS.

## Acknowledgements

We want to acknowledge the study participants and their families for taking part in this study. We thank the fieldworkers and nurses who helped with the collection of the study samples. We are also grateful for the funding of this project by the Medical Research Council, UK.

## Authors’ contributions

EFN, MA and MO conceptualised this study. MO, AAO and EO collected the study samples. EFN, KJ, FC, KJ, SB and AJ carried out the microbiological analysis of the study samples. EFN performed the phylogenomic analyses and drafted the manuscript. MA provided reagents/materials and supervised the lab work. MJP provided reagents/materials and reviewed drafts of the paper. JJ carried out the data and statistical analyses. AW managed the data. MS and NFA guided the bioinformatics analyses and drafts of the paper. All authors reviewed the manuscript.

## Availability of data and material

The data presented in this manuscript will be made available upon request.

## Competing interests

The authors have no competing interests to declare.

